# Toward an understanding of the chemical ecology of alternative reproductive tactics in the bulb mite (*Rhizoglyphus robini*)

**DOI:** 10.1101/2021.06.30.450527

**Authors:** Adam N. Zeeman, Isabel M. Smallegange, Emily Burdfield Steel, Astrid T. Groot, Kathryn A. Stewart

## Abstract

**Background:** Under strong sexual selection, certain species evolve distinct intrasexual, alternative reproductive tactics (ARTs). In many cases, ARTs can be viewed as environmentally-cued threshold traits, such that ARTs coexist if their relative fitness alternates over the environmental cue gradient. Surprisingly, the chemical ecology of ARTs has been underexplored in this context. To our knowledge, no prior study has directly quantified pheromone production for ARTs in a male-polymorphic species. Here, we used the bulb mite—in which males are either armed fighters that kill conspecifics, or unarmed scramblers—as a model system to gain insight into the role of pheromones in the evolutionary maintenance of ARTs. Given that scramblers forgo investment into weaponry, we tested whether scramblers produce higher pheromone quantities than fighters, which would improve the fitness of the scrambler phenotype, e.g. through female mimicry to avoid aggression from competitors. To this end, we sampled mites from a rich and a poor nutritional environment and quantified their production of the female sex pheromone α-acaridial through gas chromatography analysis.

**Results:** We found a positive relationship between pheromone production and body size, but males exhibited a steeper slope in pheromone production with increasing size than females. Females exhibited a higher average pheromone production than males. We found no significant difference in slope of pheromone production over body size between fighters and scramblers. However, scramblers reached larger body sizes and higher pheromone production than fighters, providing some evidence for a potential female mimic strategy adopted by large scramblers. Pheromone production was significantly higher in mites from the rich nutritional environment than the poor environment.

**Conclusion:** Further elucidation of pheromone functionality in bulb mites, and additional inter-and intrasexual comparisons of pheromone profiles are needed to determine if the observed intersexual and intrasexual differences in pheromone production are adaptive, if they are a by-product of allometric scaling, or diet-mediated pheromone production under weak selection. We argue chemical ecology offers a novel perspective for research on ARTs and other complex life-history traits.

## BACKGROUND

Animals exhibit various behavioral, morphological and physiological adaptations related to their ability to attract and compete for mates [1]. Many of these adaptations have evolved under strong sexual selection, resulting in distinct, intrasexual Alternative Reproductive Tactics, or ARTs [2]. ARTs are pervasive in nature [e.g. 3-8] and explanations for their maintenance include models of frequency-dependent selection on genetic polymorphisms (i.e. the fitness of each ART depends on the relative frequency of all ARTs in a population; 9) and models of condition-dependent developmental divergence [4, 10-12]. Predominantly, however, ART expressions are considered threshold traits, wherein one tactic is expressed when an environmental or (environmentally-driven) physiological cue reaches a certain, genetically determined threshold during ontogeny, and the alternative tactic is expressed if this threshold is not reached [e.g. 13, 14; see also 2].

Importantly, such environment-dependent ARTs can only coexist if the fitness functions of the ARTs cross over the gradient of the environmental cue [15]. A salient cue of ART expression is diet [14, 16, 17], because nutritional quality and quantity dictate the body sizes and resource budgets of developing individuals [18], thereby affecting the potential future mating success of different ARTs [2]. For example, large juvenile males likely have sufficient resources to develop into large adults with morphological structures that can be used as weapons when defending their mate against rival males, whereas small juvenile males are unlikely to be successful during combat, and most likely benefit more by refraining from developing weapons to resort to “sneaking” tactics [19]. Such sneaking tactics have been observed across a wide range of taxa [e.g. 4, 7, 20], and would be particularly successful if small(er) males can additionally conceal themselves from rival, fighter males, for example by mimicking a female [21].

Female-mimic mating strategies are found in a variety of taxa [e.g., ruffs, 22; snakes, 23; marine isopod, 24]. Sometimes mimicking mechanisms are visual [25], but female mimicry also exists in organisms that rely on non-visual cues [e.g., chemical/pheromone cues, 24]. In many animal taxa, pheromones are the primary means of intraspecific communication and mate attraction [26-28]. Communication through pheromones is subject to strong sexual selection as its effectiveness influences the fitness of both the signaler and receiver [29]. Despite these prominent selective forces however, pheromone profiles can vary strongly between individuals of the same species—both in quantity and composition [30]. Indeed, this variation in an individual’s pheromone profile is often mediated by environmental factors [31, 32], especially diet [33-40].

For example, diet has been demonstrated to influence pheromone profiles directly via essential pheromone precursors [39], and indirectly, through an increased resource allocation towards pheromone production [36]. In fact, within heterogeneous environments, where food availability is highly variable between individuals, strong sexual selection may act on divergent diet-mediated pheromone profiles and drive the evolution of different pheromone-based ARTs. For example, well-fed individuals may attempt to attract mates by producing high quantities of sex pheromone, while poorly fed individuals may attempt to be inconspicuous by producing low pheromone quantities, permitting them to sneak past competitors to gain access to females.

Although inconspicuous sneaking, or female mimicry, tactics are known to occur in many species with male ARTs [e.g., 4, 5, 7], in most cases, the role of pheromones in these strategies remains unknown [but see 24].

The aim of this study was to gain insight into the importance of pheromones in the maintenance of ARTs. For this, we used the bulb mite (*Rhizoglyphus robini*), a well-studied example of a system in which ART expression is a threshold trait cued by diet. Importantly, ART expression in *R. robini* does not depend on population density, unlike in its sister species *R. echinopus* [41], nor on ART frequency [42]. Upon maturity, male *R. robini* develop into one of two distinct morphs (see Figure 1): (1) juvenile males that are relatively large mostly mature as “fighters”, which possess a thickened third leg pair with a sharp end that can be used to kill conspecifics [45, 46], and (2) juvenile males that are relatively small mostly mature as “scramblers”, which lack the weaponized leg pair (although a rare third morph, the mega-scrambler, has recently been suggested; [47]). Although scrambler expression is regulated by a (partially) genetically determined threshold for body size [17; 48; 49], gene-by-environment interactions also play a key role [50], with diet quality and quantity thought to be the primary drivers of body size and therefore ART expression [18]. Despite this, the selection pressures that maintain the coexistence of fighters and scramblers are still not fully understood. Numerous studies have attempted to identify or quantify the fitness functions that underlie this evolutionary maintenance [17, 42, 44, 48, 50-52], but some facets of bulb mite ecology, such as its chemical communication, remain unexplored in this context.

**Figure 1:**
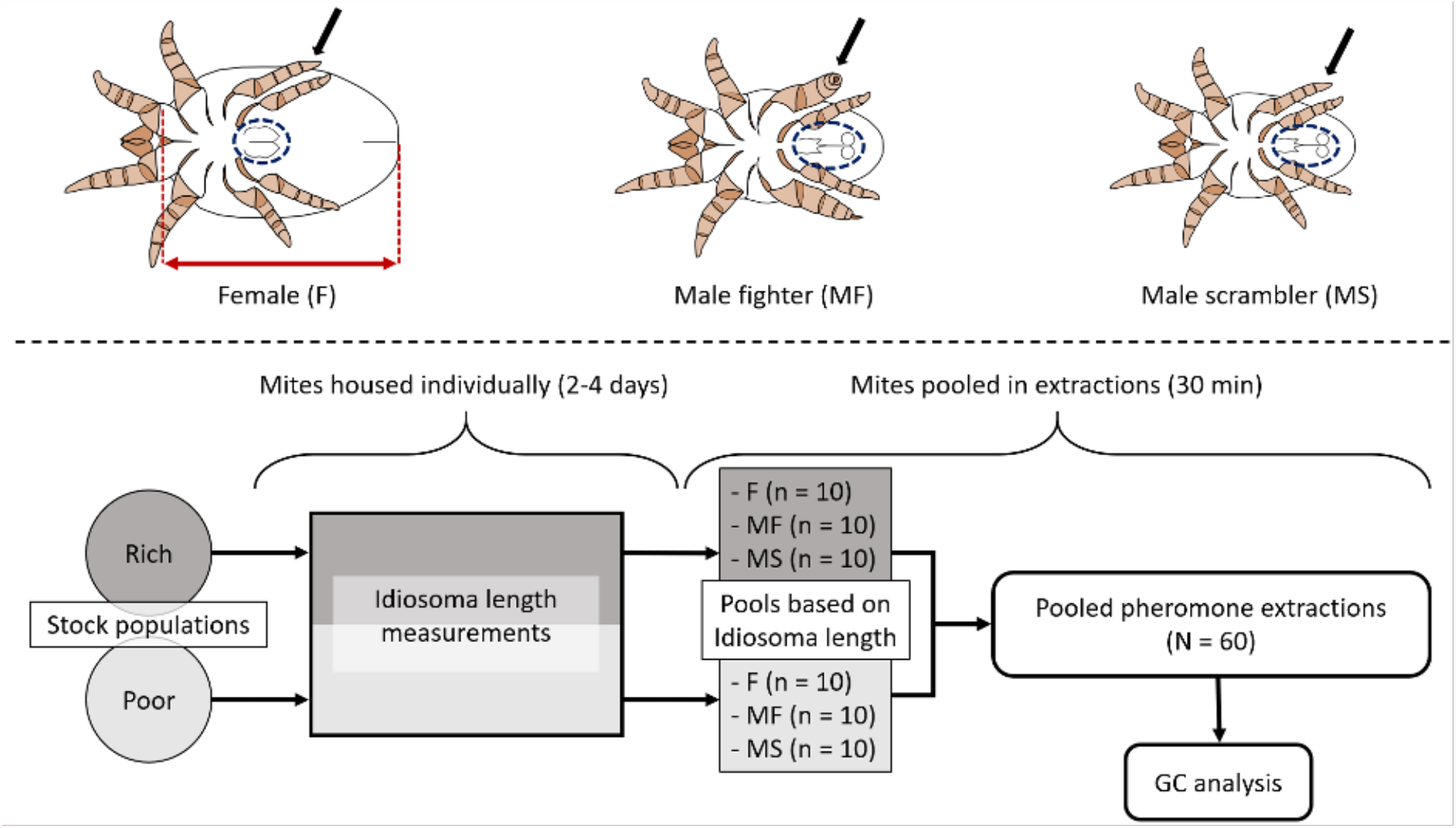
A) Ventral drawings of adult bulb mites (*Rhizoglyphus robini*). From left to right: emale, male fighter and male scrambler. Morphological characteristics useful for identification are highlighted: black arrows indicate the third leg pair (thickened and sharply terminated in fighters), and dotted blue ellipses highlight the genitalia (including the anal discs in the males), which differ distinctly between males and females [43]. The idiosoma length is indicated with a red arrow (between the two dashed red lines; 43, 44) on the drawing of the female. Mite drawings by F. Rhebergen. B) Workflow for pheromone quantifications: (i) females, male fighters and male scramblers were randomly selected from rich (dark grey) and poor (light grey) stock populations, and housed individually in plastic tubes, (ii) the idiosoma length of the collected mites was measured as a proxy for body size, (iii) mites from the same stock population and of the same sex or ART, were pooled based on idiosoma length, totaling 60 pools, and (iv) the α-acaridial production of each pool was quantified by performing hexane extractions followed by gas chromatography (GC) analysis.

As bulb mites are blind, we would expect them to rely heavily on chemical signals for communication [53]. Currently, a female sex-pheromone, α-acaridial [54, 55], has been identified in this species, which can elicit increased mounting behaviour in male bulb mites [55], but its role in the maintenance of fighter-scrambler coexistence is unexplored. There is also notable intrasexual variation in α-acaridial production in bulb mites males—more so than in females [55], potentially highlighting differential ART investment into pheromone production. For example, because scramblers forgo the development of weaponized legs and spend less energy on aggressive behavior compared to fighters [46], they might invest more into pheromone production. Increased pheromone production may also serve to mimic the pheromone profile of the much larger females, ultimately reducing the conspicuousness of scramblers towards competitors and allowing them to avoid costly intrasexual combat. As such, pheromones may be an important factor in the maintenance of ARTs.

Here, we conducted a laboratory experiment to test the hypothesis that bulb mite scramblers produce higher quantities of the female sex pheromone (α-acaridial) than fighters, potentially as a means of avoiding intrasexual reproductive competition. Because nutritional uptake has been shown to be an important driver of ART expression in bulb mites [17,18], and because diet strongly influences pheromone production across many invertebrates [reviewed by 56], we measured α-acaridial production of mites raised on two diets of different food quality. Also, we included body size as a covariate to control for any body size effects on α-acaridial production. Our results show that (i) α-acaridial production was positively correlated with body size, and this relationship was steeper in males than females, (ii) α-acaridial production was influenced by the nutritional environment, (iii) on average, females produced more α-acaridial than males, and (iv) there was no significant difference in slope of α-acaridial production over body size between the male ARTs.

## RESULTS

### Detection and quantification of α-acaridial

The GC-MS analysis indicated that the pheromone α-acaridial was present in the bulb mite extracts (Figure 2). Analysis of the ion fragments from the peaks on the GC-MS profiles indicated that the compound at 22.8 min retention time was in fact α-acaridial. The presence of the molecular ion at 166 m/z further supported this [see 54]. Through the GC-analysis, α-acaridial production could be quantified for 56 of the 60 pooled extracts (one outlier was excluded, resulting in N = 55). On average, females produced (mean ± SD) 13.53 ± 8.04 ng of α-acaridial (based on per-mite-averages for each pool, n = 19 pools), fighters produced 7.35 ± 4.58 ng (n = 18 pools) and scramblers produced 12.67 ± 12.88 ng (n = 18 pools).

**Figure 2:**
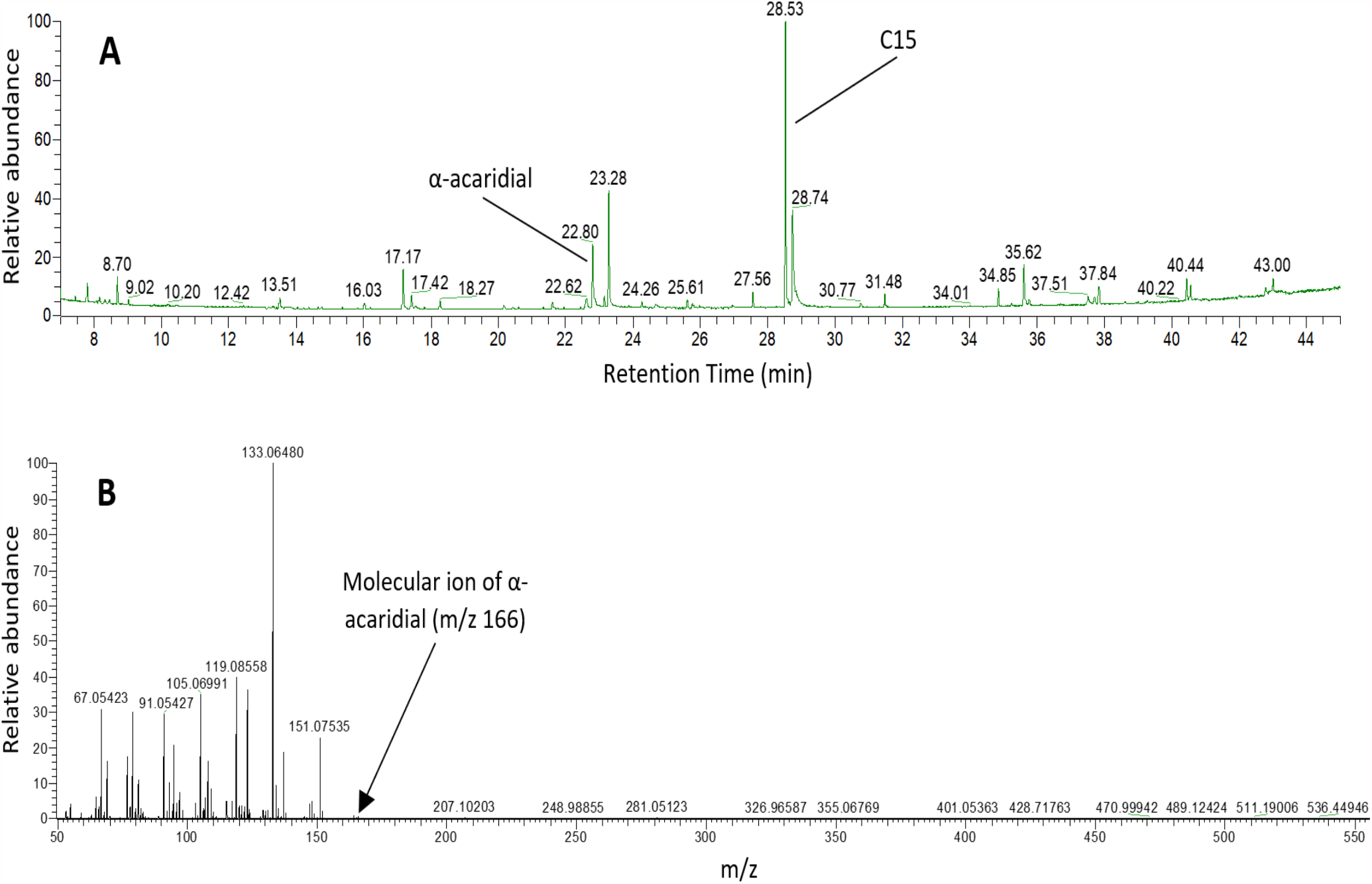
A) gas liquid chromatogram of a hexane extract from 10 *Rhizoglyphus robini* females (sampled from the poor stock populations). Numbers above peaks give their retention time. C15; pentadecane (internal standard). B) mass spectrum for the peak at 22.8 min. Numbers above peaks give their mass-to-charge ratio (m/z). The ion fragments and the presence of the molecular ion at 166 m/z indicate this compound is α-acaradial [see 54]. See methods for GC-MS conditions.

### Effects of morph, diet and idiosoma length on α-acaridial production

The model selection procedure revealed that the effects of morph, diet and idiosoma length on log(α-acaridial production) were best described by model 7 (see Table 1), in which the relationship between idiosoma length and log(α-acaridial production) differed significantly between the sexes (idiosoma length × morph; *F*_1,50_ = 15.12, *p* < 0.001), with males showing a bigger increase in log(α-acaridial production) with increasing idiosoma length than females (Figure 3a) (note that in model 7, fighters and scramblers are merged into one factor level ‘males’). Additionally, females produced on average more α-acaridial than males (*F*_1,50_ = 14.33, *p* < 0.001). Also, log(α-acaridial production) was significantly higher on the rich diet than on the poor diet (rich 2.60 ± 0.66 ng, n = 29 pools; poor 1.45 ± 0.76 ng, n = 26 pools; *F*_1,50_ = 9.65, *p* < 0.01; Figure 4).

**Table 1:**
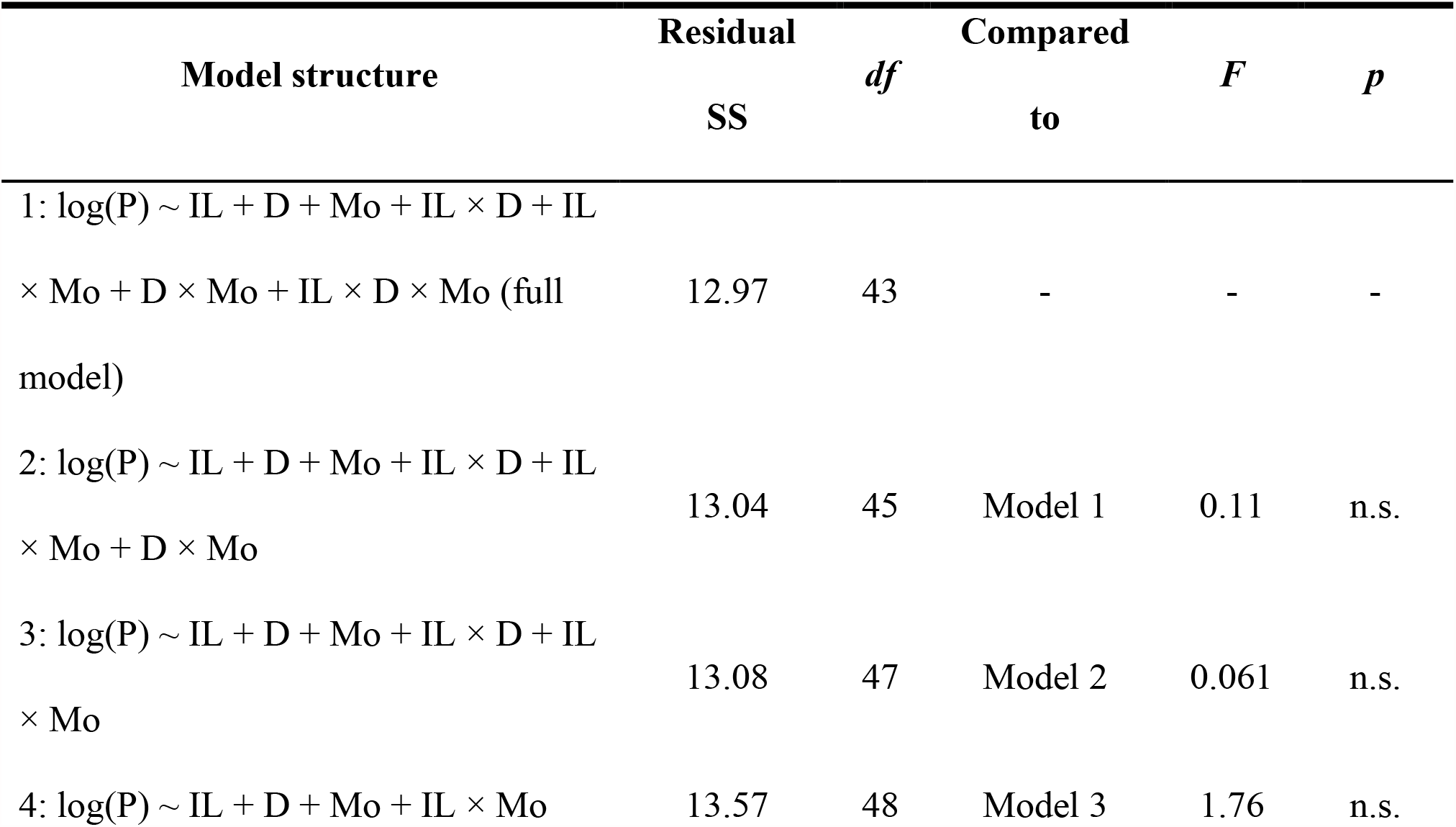
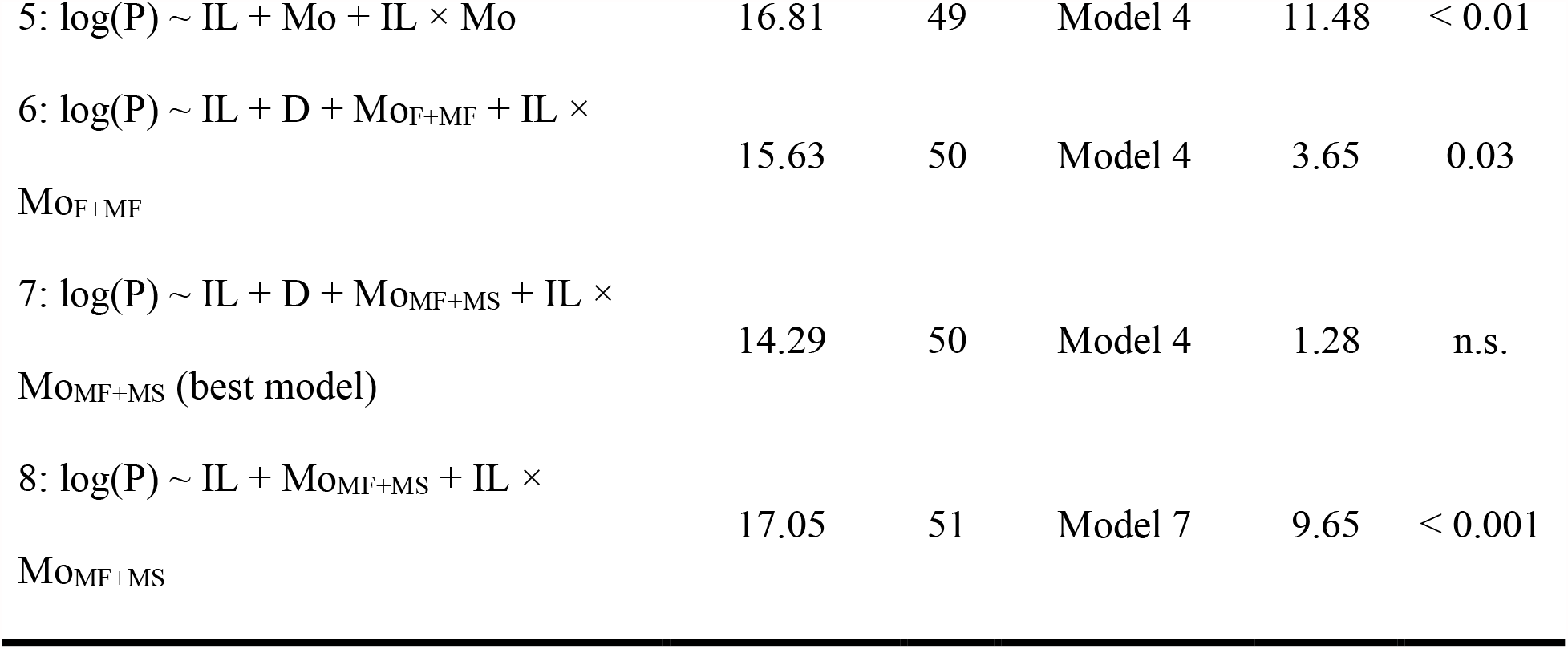
Structure and comparison of linear models where Mo is the fixed factor morph (with levels female, fighter, scrambler), D is the fixed factor diet (with levels rich, poor), IL is idiosoma length (µm), and P is the response variable log(pheromone production) (log ng) in pooled bulb mite extracts (N = 55). In model 6, females (F) and fighters (MF) were grouped together as one factor level in the variable Mo. In models 7 and 8, fighters and scramblers (MS) were grouped together. Model selection (see main text) was performed using the F-ratio test: *F* = [(*RSS*_*0*_ − *RSS*_*1*_)/(*df*_0_ − *df*_*1*_)]/(*RSS*_*1*_/*df*_*1*_), where *RSSi* and *dfi* are the residual sum of squares and degrees of freedom, respectively, of model *i* (where *i* = 0 as the simpler model and *I* = 1 as the more complex model). Non-significant (n.s.) *p*-values (> 0.05) indicate that the simpler model did significantly reduce the residual deviance compared to the more complex model.

**Figure 3:**
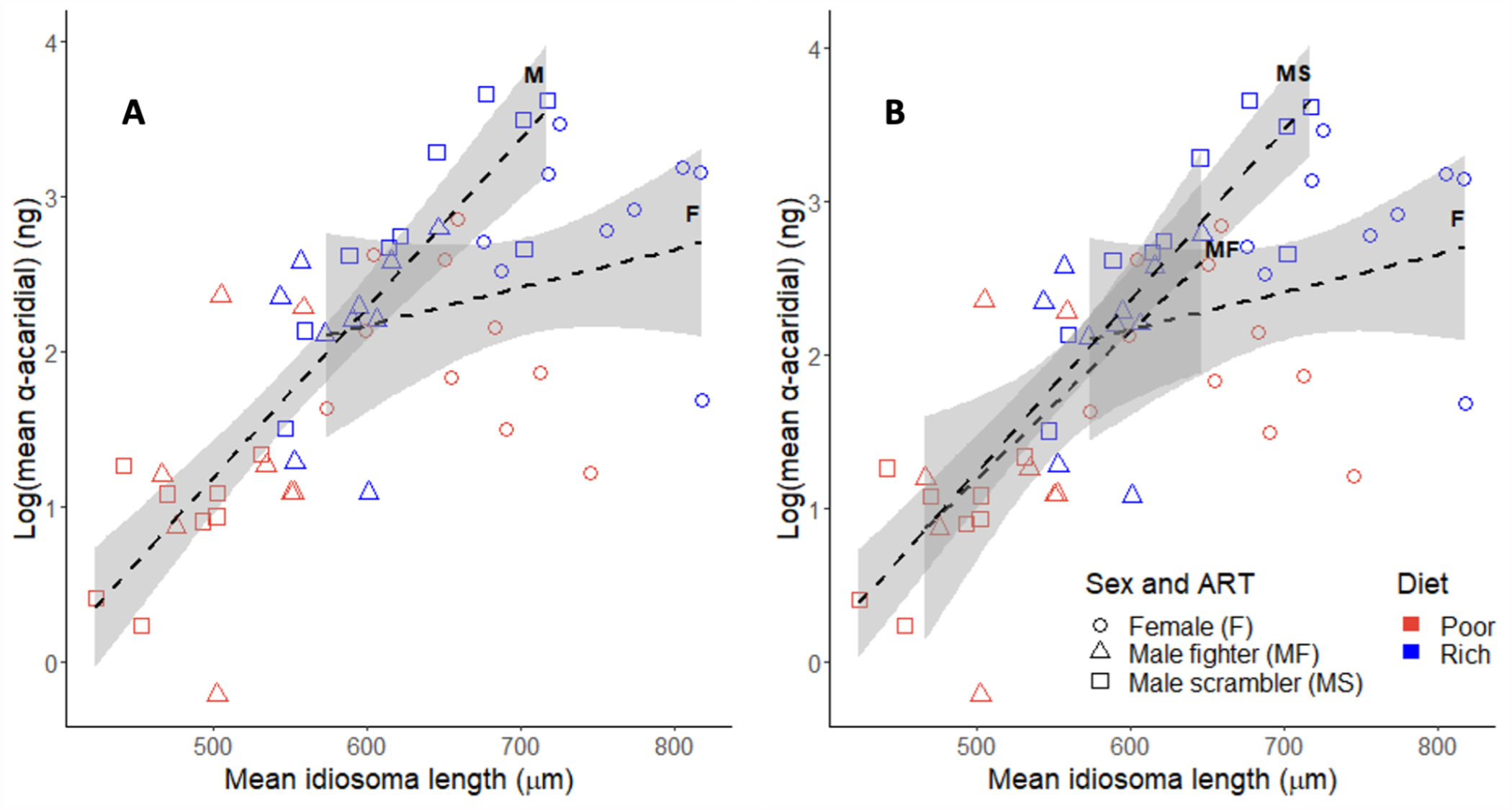
Relationships between idiosoma length and log(α-acaridial production) (shown as per-mite-averages for 55 pools) for bulb mites fed on a rich or poor diet. A) illustrated male ARTs grouped together; trendlines for females (F; n = 19) and males (M; n = 36). B) shows separate trendlines for fighters (n = 18) and scramblers (n = 18) in addition to females. Standard errors are indicated by grey shading. Legend is shown on the bottom right.

**Figure 4:**
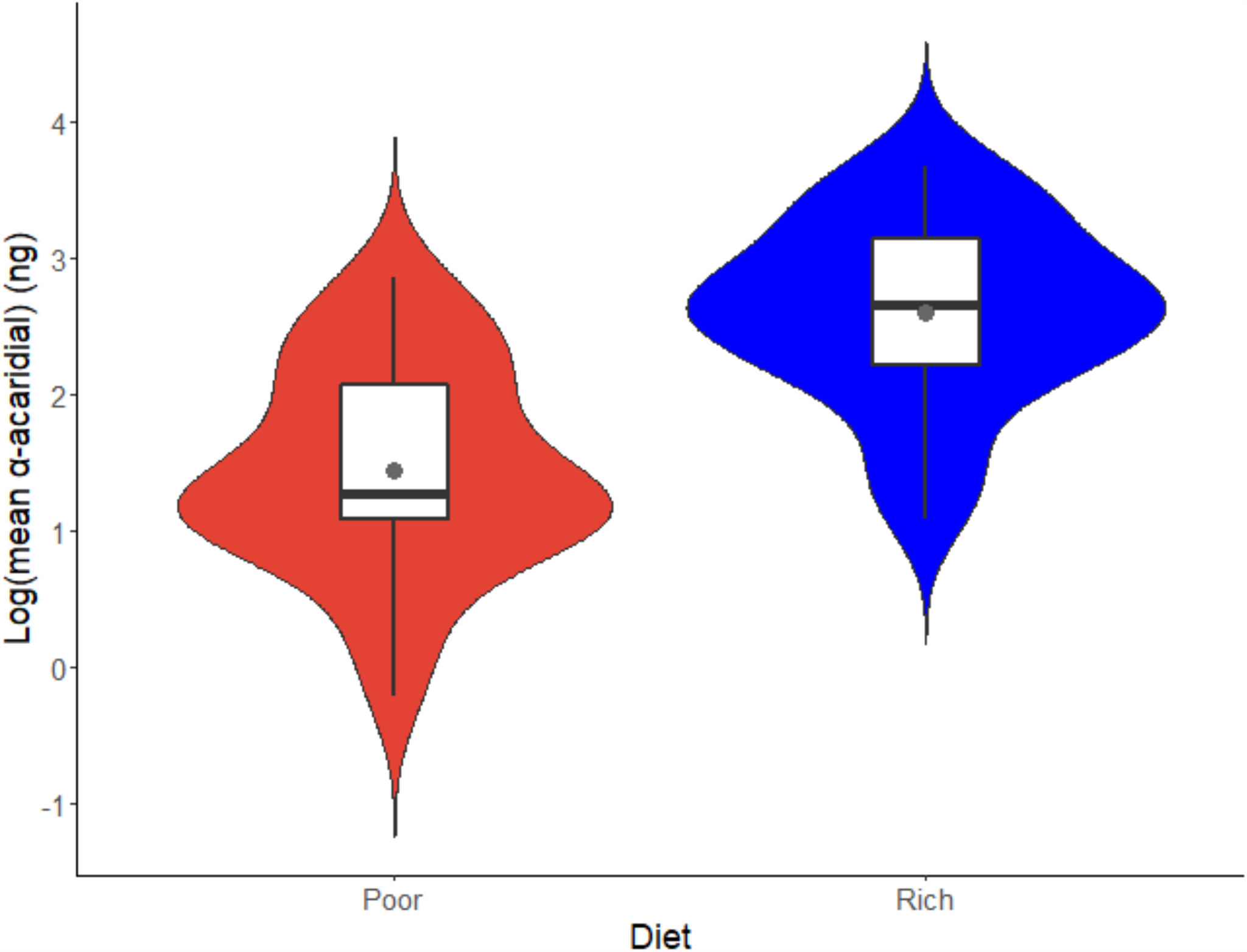
Violin plot, shown as per-mite-averages for 55 pools, of log(α-acaridial production) across nutritional environments for bulb mites fed on a rich (N = 29) or poor (N = 26) diet. Grey dot shows the group mean, black horizontal bar denotes median, boxes represent the interquartile ranges and whiskers show minima and maxima.

Inspection of the second-best fitting model (Table 1, model 4), in which fighters and scramblers were included as separate factor levels in the factor morph, revealed, like in the best-fitting model, that males of both morphs showed a bigger increase in log(α-acaridial production) with increasing size than females (idiosoma length × morph; *F*_2,48_ = 8.34, *p* < 0.001) (Figure 3b). However, inspection of this second-best fitting model also shows that the highest log(α-acaridial production) levels are associated with scramblers, which were also of longer idiosoma length than fighters. Finally, overall, mites from the poor diet populations were significantly shorter in idiosoma length than mites from the rich diet populations (rich 654.43 ± 86.55 μm, n = 316; poor 551.65 ± 93.55 μm, n = 282; two sample t-test, *t*_596_ = -13.95, *p* < 0.001).

## DISCUSSION

In our study, we aimed to gain insight into the role of pheromones in the evolution and maintenance of ARTs by assessing α-acaridial production in the male-dimorphic bulb mite under two nutritional regimes. We tested the hypothesis that scramblers, given that they forgo investment into weaponized legs, can produce higher quantities of α-acaridial than fighters, which would improve the reproductive success of the scrambler phenotype, e.g., through female chemical mimicry. We found that α-acaridial production was positively correlated with body size, and the slope of α-acaridial production over body size was steeper in males than females. In addition, α-acaridial production was influenced by the nutritional environment, as mites on the rich diet produced more pheromone than those on the poor diet. On average, females produced more α-acaridial than males. Differences between male ARTs were not incorporated in the linear model that best described the data (although they were incorporated into the second-best model), indicating that there is no significant difference in slope of α-acaridial production over body size between the male morphs. Nevertheless, scramblers reached larger maximum body sizes and higher maximum α-acaridial production compared to fighters.

### Intersexual and intrasexual differences in pheromone production

We found that females produced significantly more α-acaridial than males, corroborating the intersexual differences in α-acaridial production described by Mizoguchi *et al*. [55]. Importantly, the slope of pheromone production over body size was steeper in males than females, indicating that intersexual differences in pheromone production are not due solely to size differences between the sexes. Potentially, males may disproportionately benefit from an increased α-acaridial production with increased size. If, for example, pheromone production acts as an honest indication of quality in bulb mites, sexual selection—which is particularly strong in species with male ARTs [2]—may drive well-conditioned males to produce as much pheromone as they can, without incurring high viability costs. Alternatively, the steeper slope of pheromone production over body size for males may result from differences in chemical ecology between large and small males. The largest males, which in this study are represented mostly by scramblers, produced some of the highest pheromone quantities in the dataset, while many of the smallest males in this study produced very low quantities. This provides some evidence for the hypothesized female mimicry strategy adopted by large scramblers. These large, well-conditioned scramblers may be the only males capable of producing enough pheromones to mimic female pheromone profiles, while also reaching body sizes comparable to females. As such, these scramblers may disproportionately benefit from producing high pheromone quantities compared to smaller males. It should be emphasized however, that this study does not provide evidence that all scramblers are female mimics, as the model that best describes the data did not differentiate between male ARTs. Nevertheless, the results of this study do warrant further exploration of a female mimic or “sneaking” strategy in bulb mites, particularly because ‘mega-scramblers’—which sometimes elicit mating behavior from other males—are suggested to be a third ART [47]. Indeed, these mega-scramblers may be the result of sexual selection driving larger scramblers to chemically and physically resemble females. Intriguingly, some of the largest scramblers in this study, which produced higher pheromone quantities than most females in the dataset, exceeded a body size breakpoint for mega-scramblers calculated by Stewart *et al*. [47], implying these individuals may in fact represent the mega-scrambler trimorphism.

### Role of nutritional environment and body size in pheromone production

The observed effects of body size and diet both suggest that pheromone production is linked to nutritional uptake—particularly because body size was also dependent on the nutritional environment. The positive correlation between body size and pheromone production may stem from covariation of these variables with diet quality and uptake. As such, individuals that can consume more high-quality food subsequently grow larger [17], while simultaneously maintaining a good enough condition to produce high pheromone quantities. In this context, pheromone production could be a form of honest signaling of individual quality [56]. Alternatively, larger individuals may simply produce more pheromone because they have bigger or better developed pheromone glands, and as such, the observed patterns in pheromone production may be non-adaptive. Indeed, there is some support for allometric scaling of the production of defensive chemicals in astigmatic mites, as a study on *Archegozetes longisetosus* showed that the quantity of defensive chemicals from opisthonotal glands scales allometrically with body mass during ontogeny [57]. However, it is not known if this allometric relationship also holds for adult mites with variable mass or body size. Studies on the relationship between body size and pheromone production in other invertebrates have yielded inconsistent results, as positive correlations were found in some species [58-61] while correlations were absent in others [35, 62, but see 58]. Regardless of what drives the observed correlation between body size and pheromone production, the results of this study provide further evidence that the nutritional environment is an important driver of pheromone production, at least in bulb mites. This provides some support for the hypothesis that pheromones are energetically demanding to produce in this species. High nutritional intake likely allows for increased allocation of resources towards pheromone production, which is common among invertebrates [reviewed by 56]. And while it remains unclear to date how sex pheromones in arthropods are largely biosynthesized, evidence suggests acquisition through diet or endosymbionts rather than *de novo* [e.g., 63], further aligning with the body size and nutritional environment patterns shown here.

### From the chemical ecology of bulb mites to the evolution and maintenance of ARTs; current knowledge and future directions

To better understand the ecological significance of divergent pheromone profiles between sexes and (potentially) male ARTs in bulb mites, it is imperative that pheromone functionality is further elucidated in this species. So far, functionality has only been described for two bulb mite exudates; α-acaridial as a putative female sex pheromone [55] and neryl formate as an alarm pheromone [64]. As a putative female sex pheromone, α-acaridial was a good candidate to investigate the role of pheromones in the maintenance of bulb mite ARTs. However, evidence for the sex pheromone activity of α-acaridial is limited to a single study, where the compound was found to be present in the fractions of female hexane extracts that triggered mounting behavior in males [55]. Synthetic α-acaridial was also shown to elicit mounting behavior at a dose of 10 ng. However, neither of these findings prove that α-acaridial is the only female sex pheromone in bulb mites. Indeed, it is possible that various compounds function synergistically (with or without α-acaridial) as a sex pheromone. These synergistic interactions are less likely to be detected in bioassays with separated fractions of bulb mite extracts. Furthermore, the fact that α-acaridial was also found in males, at quantities that greatly exceed the active dose ([55] reported an average quantity of 163 ± 97 ng for males), suggests that the functionality of this compound as a sex pheromone may be reduced or enhanced by other compounds. Such synergistic or antagonistic pheromone effects are well known in pheromone signaling systems [e.g. 65-68; see also 27], and should always be considered when assessing the effects of individual compounds.

Other bulb mite exudates with potential relevance to the chemical ecology of this species include neral [64], several hydrocarbons [69)] robinal, perillene and isopiperitenone [70]. Several of these compounds have also been found in other astigmatic mites [71], but their functionality is mostly unknown in these species.

The lack of knowledge on bulb mite chemical ecology means that there are many avenues for research into pheromone functionality in this species to determine if the intersexual and (to a lesser extent) intrasexual patterns in pheromone production, observed here and by Mizoguchi *et al*. [55], are adaptive or merely a byproduct of allometric scaling or other factors. This would help us to resolve if large females and scramblers produce high pheromone quantities to gain a fitness advantage or because they have larger pheromone glands. Pheromone functions could be elucidated by exposing mites in various ecological settings to the different compounds found in bulb mite extracts [54, 64, 69, 70], including different combinations and concentrations of these compounds to test for synergistic or antagonistic effects. Given the apparent importance of diet in pheromone production observed in this study, further research on how the nutritional environment mediates (e.g., via biosynthesis) pheromone production in bulb mites is warranted. For example, it is unknown to what extent adult pheromone production is driven by nutritional uptake before maturation. In some mite species, juveniles lose the contents of their glands during each molt [72], suggesting that they may be incapable of storing pheromones or other chemicals through their development. The importance of juvenile and adult diets for pheromone production can be investigated by maintaining developing mites on rich and poor diets and switching the diet at maturity for some treatment groups. Similar studies have been conducted in other invertebrates; Jensen *et al*. [40] showed that a rich adult diet can compensate for the effects of a poor diet during ontogeny in male cockroaches, while Edde *et al*. [35)]found that adult diet, but not diet during the larval stage, affected pheromone production in the beetle *Rhyzopertha dominica*. Thus, our study should represent a springboard for myriad future investigations into the role of chemical ecology on the maintenance of ARTs.

In fact, extending beyond mites, there are other promising model systems for studying the link between chemical ecology and ART maintenance. For example, in some male-polymorphic blennies (*Salaria pavo* and *Salaria fluviatilis*), sneaking fertilization occurs from female-like male ARTs that lack anal glands and thus a putative sex pheromone [73]. In the black goby, in which males are large “parentals”, small sneakers or an intermediate phenotype [74], males produce a sex pheromone that triggers aggression in other males [75]. However, parental males were found to react aggressively to the pheromone-containing ejaculate of other parentals but not to the ejaculate of sneakers, indicating that sneaker males are pheromonally inconspicuous. The examples outlined above suggest that pheromones likely play a key role in the success of ARTs in many species, especially when considering the prominent role of pheromones in intraspecific (sexual) communication throughout nature [26-28]. Therefore, future research on ARTs would benefit from comparing pheromone profiles between male ARTs and females (i.e., to explore pheromone based female mimicry) in a variety of taxa.

The direction of putative evolutionary relationships between ARTs and within-population variation in pheromone profiles is another area bearing investigation. So far, we have briefly speculated on how sexual selection may act disruptively on pheromone profiles in heterogeneous environments, and how this may promote the evolution of different pheromone-based ARTs.

However, evidence for disruptive selection on pheromone profiles in natural populations is rare [76]. Instead, pheromone profiles and other forms of sexual communication are often under stabilizing selection [77-80]. Therefore, the evolutionary relationship between ARTs and divergent pheromone profiles may be reversed, such that the evolution of ARTs *facilitates* the evolution of divergent pheromone profiles. By definition, ARTs adopt different strategies to improve their reproductive output, and therefore they face different selection pressures [2]. For example, large males that compete for females directly will likely be favored by selection to develop traits that improve their ability to fend off competitors, while smaller males that adopt sneaking tactics may well be favored to develop traits that make them inconspicuous towards other males. These divergent selective pressures may decouple male (sex) pheromone profiles in the population from stabilizing selection, or rather, stabilizing or directional selection may now occur more or less independently for the pheromone profiles of both ARTs, leading to disruptive selection on the population level. There is also emerging evidence that variation in sex pheromone profiles can be maintained by balancing selection [81, 82], e.g., through heterozygote advantage [81]. Thus, within-population variation in pheromone profiles may arise and be maintained through various mechanisms. Further research on the chemical ecology of species with ARTs is needed to assess if divergent pheromone profiles within populations facilitate the evolution of ARTs or vice versa.

## CONCLUSIONS

We found a positive relationship between pheromone (α-acaridial) production and body size in bulb mites, but importantly, males demonstrated a steeper slope in pheromone production with increasing size than females. We found no significant difference in slope of pheromone production over body size between fighters and scramblers, but scramblers reached larger maximum body sizes and thus had higher maximum pheromone production compared to fighters. The results of this study also indicate that diet quality influences pheromone production in bulb mites, further highlighting the importance of the nutritional environment for several aspects of the ecology of species displaying environmentally-cued ARTs. The observed patterns of intersexual and intrasexual differences in pheromone production may be adaptive, as sexual selection may have driven the evolution of divergent pheromone profiles that relate to different, condition-dependent strategies, such as sneaking in males. The observed patterns may also be non-adaptive however, potentially reflecting allometric scaling of pheromone production with body size, or diet-mediated pheromone production under weak selection. Further elucidation of pheromone functionality in bulb mites, and additional inter-and intrasexual comparisons of pheromone profiles, are needed to assess the role of pheromones in the maintenance of male-polymorphism in this species.

To our knowledge, this is the first study to directly quantify the production of a pheromone for two ARTs in a male polymorphic species. Yet, intrasexual differences in pheromone production in male-polymorphic species offer promising research avenues in the context of crossing fitness functions that underlie the maintenance of these polymorphisms. Importantly, a more complete understanding of complex life-history traits, such as ARTs, requires investigation through interdisciplinary contexts, such as eco-evolutionary dynamics, developmental biology, population genetics and indeed, chemical ecology.

## METHODS

### The bulb mite

The blind bulb mite (*Rhizoglyphus robini*), a common agricultural pest that feeds on various crops [83], is an excellent model system for studying the expression and maintenance of ARTs. In addition to its short generation time and high reproductive output, this microscopic mite can easily be reared in the laboratory under various conditions [53]. After hatching, bulb mites undergo four or five developmental stages: larva, protonymph, deutonymph (a facultative dispersal stage that occurs under adverse conditions, such as food or water scarcity), tritonymph and adult [84]. Transitions between these stages occur in the form of a quiescent molting stage. Upon maturity and depending on the nutritional environment, male bulb mites develop into either armed fighters, or benign scramblers (Figure 1).

### Maintenance of stock populations

Stock cultures originated from flower fields near Anna Paulowna (The Netherlands), where they were collected from flower bulbs in 2010. Up until the COVID-19 outbreak in March 2020, stock cultures were kept in an unlit climate chamber (25 ± 1 °C, 60% relative humidity) at the Institute for Biodiversity and Ecosystem Dynamics at the University of Amsterdam (The Netherlands). After that, they were moved to a location without access to climate chambers and kept at room temperature. The experiments were conducted at room temperature as well. Mites were kept in sealed but ventilated plastic containers (50 mm high, 85 mm in diameter) that contained a layer of plaster of Paris (∼15 mm thick) that was nearly saturated with water. The stock cultures were either always given *ad lib* access to dried yeast granules (Bruggeman instant yeast), or *ad lib* access to grains of rolled oats. Yeast and oats are of high and low nutritional quality, respectively, due to their respective high and low protein content [50]. Therefore, these resources will further be referred to as “rich” (yeast) and “poor” (oats) diets. To reduce inbreeding, stock populations fed on the same diet were intermixed periodically, effectively creating multiple meta-populations. Additional food and water were provided to each stock container once or twice per week (in a manner similar to [44]). The observed heterozygosity (averaged across sex and ART) of the stock populations was measured at 0.39 for the rich environment and 0.48 for the poor environment (50).

### Experimental setup

#### Sampling stock mites

The mites used in the experimental procedures were all randomly selected from the stock cultures (Figure 1). The mites were handled (using a fine brush or a metal probe) and identified under a ZEISS Stemi 508 stereomicroscope; their life-history stage, sex and ART were determined based on their body size, genitalia and on the morphology of their third leg pair (Figure 1). Mega-scramblers were not used in any of the experimental procedures due to their rarity within the stock populations [47]. After collection, mites were housed individually (to avoid cannibalism and mating) in sealed, ventilated plastic tubes (50 mm high, 16 mm in diameter) that were filled up to three-fourths with a mix of plaster of Paris and charcoal powder for visual contrast. The plaster mix was made in batches by adding 40 mL tap water to 40 g plaster powder and one-third of a tablespoon of charcoal powder. The plaster was left to harden in the tubes for at least 24 hours at room temperature before the tubes were used. The tubes were closed off with a cap containing a small air hole and a piece of fine mesh to prevent escaping while still allowing air flow. Each tube was hydrated—before the mites were placed inside—by adding two drops of tap water on the dried plaster layer with a drip pipette. After the mites were collected, one yeast granule was added to the tubes that housed mites from the rich stock populations, and one-fourth of an oat grain was added to the tubes that housed mites from the poor stock populations. Food was added in limited quantities to prevent mold growth on uneaten leftovers. All collected mites were kept in their individual tubes until they were used in pooled pheromone extractions (see below for details)—usually this constituted a period of 2-4 days in the tubes.

#### Body size measurements

The idiosoma length (the length of the body without mouthparts; Figure 1D) of the collected mites was measured as a proxy for body size [43, 44]. First, the mites were photographed using a ZEISS Axiocam 105 color camera at 0.63-5 × magnification that was connected to a Zeiss Stemi 2000-C stereomicroscope. From these photos, the idiosoma length was measured to the nearest µm using Zen lite (Blue edition) analysis software. After enough individuals were measured (see below for details), mites from the same stock population that were of the same sex or ART were pooled for pheromone extractions based on similar idiosoma length—such that the variance in idiosoma length was as low as possible within each pool. The pooling of multiple mites was done to ensure quantifiable amounts of pheromone could be extracted (see Supplementary Information). In total, 60 pools were created from measured mites (10 pools of females (n = 8-11), 10 pools of fighters (n = 10-14), and 10 pools of scramblers (n = 10-14) from the rich and the poor stock populations).

#### Pheromone extraction and gas chromatography (GC) analysis

Mites selected for pooled extractions were removed from their individual tubes and submerged together for 30 (± 1) minutes inside a screw-top glass vial filled with 50 µl of hexane containing 200 ng of pentadecane as the internal standard. The hexane extracts were then separated from the mites using a 100 µl Hamilton syringe, and stored in crimp-top vials at −20 °C until further use. All pheromone extractions were performed within 30 hours of the body size measurements. Once all extractions were completed, the extracts were prepared for gas chromatography (GC) analysis; extracts were evaporated down to 1-3 µl under a gentle nitrogen stream (at room temperature) and topped by ∼1 µl of octane to prevent further evaporation. After preparation, the extracts were injected into a splitless inlet of a HP6890 GC coupled with a high resolution polar capillary column (DB‐WAXetr [extended temperature range]; 30 m × 0.25 mm × 0.5 µm) and a flame ionization detector (FID). The extracts were analyzed in three consecutive GC runs within a span of 72 hours. Finally, α-acaridial quantities were calculated through integration of the putative α-acaridial peaks. Integration results were corrected by the differential response of the FID to the standards in each extract. Because the number of mites differed between pooled extractions, pheromone quantities of each extract were divided by the number of mites in that extract, to get the average per mite for each pool.

#### Gas chromatography-mass spectrometry (GC-MS) analysis

To confirm the presence of α-acaridial in the hexane extracts of the mites, gas chromatography-mass spectrometry (GC-MS) analysis was performed with three pooled extracts of females from poor stock populations. The extracts used for this analysis were obtained in the same manner as the extracts used for the GC analysis. The GC-MS analysis was performed using a Thermo Trace 1300 GC and Thermo Exactive Orbitrap MS (50 to 550 m/z scan range) operated at 70 eV in a splitless mode, with a DB5-MS capillary column (30 m × 0.25 mm × 0.25 µm). Helium was used as the carrier gas, and was delivered at 1.0 ml/min. The temperature was programmed to increase from 50 °C (1.5 min hold) to 320 °C at a rate of 5 °C /min.

### Statistical analysis

Using linear regression models, we analyzed the effects of the fixed factor morph (female, fighter or scrambler), fixed factor diet (rich or poor) and continuous variable idiosoma length (μm), and all their interactions, on log(α-acaridial production) (log ng) as the response variable (α-acaridial production was log-transformed because the untransformed values were right-skewed). We identified one female pool from the rich diet as an outlier (Grubbs test for one outlier, G = 4.99, U = 0.54, p < 0.001), and we had missing values resulting from unclear α-acaridial peaks for two fighter pools and two scrambler pools from the poor diet. Because of this unbalanced data structure, we used type III sums of squares. To identify significant treatment effects, we used a model simplification procedure whereby the full model was fitted, after which the least significant term was removed (starting with the highest order interaction) in case this deletion caused an insignificant increase in deviance (significance was assessed by performing a F-ratio test) (Table 1) [85]. This procedure was repeated until the model only contained significant terms (P < 0.05). It turned out that this minimal model contained the fixed factor morph, which has three levels (female, scrambler, fighter). To assess which of the different levels of the factor morph did not significantly different from each other, we merged different, pairwise combinations of the three different levels. For example, if a model where the morph levels fighter and scrambler are merged into one level ‘males’ does not result in a significant increase in residual deviance compared to the model where morph has three levels, inference is that fighters and scramblers do not significantly differ in log(α-acaridial production). We confirmed that the assumption of a Normal error distribution was justified by visual inspection of histograms of model residuals and normal quantile-quantile plots, and confirmed that the assumption of homogeneity of residuals was justified using residuals-versus-fits plots. All statistical analyses were conducted in RStudio [86].

## LIST OF ABBREVIATIONS

ART: Alternative Reproductive Tactic
FID: Flame ionization detector
GC: Gas chromatography
GC-MS: Gas chromatography-mass spectrometry

## DECLARATIONS

### Ethics approval and consent to participate

In accordance with the Dutch 2014 Animal Experiments Act (https://wetten.overheid.nl/BWBR0003081/2014-12-18) (in Dutch), no ethics approval or consent is required for experiments on mites, as regulations within the Act only apply to vertebrates and cephalopods.

### Consent for publication

Not applicable.

### Availability of data and materials

The datasets generated and/or analysed during the current study are available in the Figshare repository (uploaded upon manuscript acceptance).

## Author’s Contributions

KAS and IMS conceived of the project and assisted in data analysis and writing the paper. EBS and ATG assisted in pheromonal extractions and EBS assisted in chemical analyses. ANZ collected mite morphological measurements and pheromonal extractions, assisted in data analyses and led the writing of the paper. All authors contributed to data interpretation and edited the final version of the paper.

## Acknowledgments

We would like to thank Samira Absalah for performing most of the GC-MS analysis, Dennis van Veldhuizen and Peter Kuperus for laboratory assistance, and Flor Rhebergen for providing bulb mite drawings and helpful discussions and feedback. Finally, we want to thank Dirk and Hella Zeeman, for allowing a mite lab to be moved into their residence during the COVID-19 outbreak.

## SUPPLEMENTARY INFORMATION: pilot hexane extractions

To ensure quantifiable amounts of pheromone could be obtained from bulb mite hexane extracts, pilot extractions were performed. Individual mites and groups of mites, ranging from two to ten individuals, were submerged in various amounts of hexane for different durations (Table S1). All mites used in these extractions were randomly sampled from the stock populations, following the procedure described in the Methods. The extracts were analyzed through gas chromatography (GC), also following the procedure described in the Methods. Pilot extractions were deemed successful when clear, quantifiable pheromone peaks were seen in the resulting chromatograms. The results indicated that at least two females or ten males (mostly performed using fighters) were required to consistently obtain measurable amounts of pheromone from a single extract. Two additional pilot extractions were performed to check for potential contamination of yeast granules and oat grains (Table S1, bottom rows). This was done because food particles from the plastic tubes that housed the mites were sometimes accidentally submerged in the hexane along with the mites during the extractions. A yeast granule and an oat grain were individually submerged in 50 µl of hexane for 30 minutes. The resulting chromatographs did not contain notable peaks, indicating that these food particles would not contaminate mite extractions.

**Table S1:** Overview of the pilot extractions. The number of mites in each extraction, the sex and ART (for males), diet, amount of hexane used in the extraction and the extraction time (i.e., how long the mites were submerged in hexane) are given. The final column indicates whether the extractions resulted in quantifiable pheromone peaks.

| **Number of**<br>**mites** | **Sex and**<br>**ART** | **Diet** | **Hexane in**<br>**extract (μl)** | **Extraction**<br>**time**<br>**(minutes)** | **Clear pheromone**<br>**peak in**<br>**chromatogram** |
| --- | --- | --- | --- | --- | --- |
| 1 | Female | Rich | 10 | 3 | No |
| 1 | Female | Rich | 50 | 10 | No |
| 8 | Female | Rich | 50 | 10 | Yes |
| 10 | Female | Poor | 50 | 10 | Yes |
| 4 | Female | Rich | 50 | 30 | Yes |
| 3 | Female | Rich | 50 | 30 | Yes |
| 2 | Female | Rich | 50 | 30 | Yes |
| 1 | Female | Rich | 50 | 30 | No |
| 1 | Fighter | Rich | 50 | 30 | No |
| 2 | Fighter | Rich | 50 | 30 | No |
| 3 | Fighter | Rich | 50 | 30 | No |
| 3 | Scramble<br>r | Poor | 50 | 30 | No |
| 5 | Fighter | Rich | 50 | 30 | No |
| 10 | Fighter | Rich | 50 | 30 | Yes |
| Yeast granule |  |  | 50 | 30 | No |
| Oat grain |  |  | 50 | 30 | No |

